# Infective prey leads to a partial role reversal in a predator-prey interaction

**DOI:** 10.1101/2021.03.15.435444

**Authors:** Veijo Kaitala, Mikko Koivu-Jolma, Jouni Laakso

## Abstract

An infective prey has the potential to infect, kill and consume its predator. Such a prey-predator relationship fundamentally differs from the classical Lotka-Volterra predator-prey premise because the prey can directly profit from the predator as a growth resource. Here we present a population dynamics model of partial role reversal in the predator-prey interaction. We parametrize the model to represent the predator-prey interaction of sea cucumber *Apostichopus japonicus* and bacterium *Vibrio splendidus*. We observe that two major factors stabilize the predator-prey interaction. First, the partial role reversal in the predator-prey community stabilizes the predator-prey interaction. Second, if the predator is a generalist and follows the type I functional response in attacking the prey, the predator-prey interaction is stable. We also analysed the conditions for species extinction. The extinction of the prey, *V. splendidus*, may occur when its growth rate is low, or in the absence of infectivity. The extinction of the predator, *A. japonicus*, may follow if either the infectivity of the prey is high or a moderately infective prey is abundant. We conclude that partial role reversal is an underestimated subject in predator-prey studies.

## Introduction

The ability of a prey to utilize the predator as a food source is referred to as a role reversal in predator-prey interaction [1–3]. The prey may become an enemy to the predator. If the role reversal is not complete the predator continues to hunt the prey while becoming vulnerable to the predation itself. In the partial role reversal the growth of the prey population relies on the prey’s normal growth rate and on the additional resource acquiring by the infectivity, in particular, by its efficiency in killing and converting the predator into nutrition.

A partial role reversal in the aquatic environment can occur in the aquaculture of sea cucumbers *(Apostichopus japonicus)* which feeds bottom sediments inhabited by the opportunistic, potentially infective bacterium *Vibrio splendidus*. The sea cucumber belongs to the class Holothuroidea in the Phylum Echinodermata. It is a bottom dwelling marine deposit feeder that uses its tentacled mouth to consume the topmost sediment layer [4,5]. The sediment contains plant and animal debris, protozoa, diatoms and a diverse selection of bacteria [6–10]. The sediment also hosts the bacteria *V. splendidus* [10,11] which has been associated with seasonal epidemics of high mortality among the cultured sea cucumbers [12,13]. On the other hand, *V. splendidus* can also coexist in the gut of healthy sea cucumbers [14,15]. Because bacteria generally form an important food source for the sea cucumber, *A. japonicus* [5] can be treated as a predator to *V. splendidus*. The interaction is not tight in the sense of traditional Lotka-Volterra predator interaction since both species can also consume other resources.

We address the problem of partial role reversal in the predator-prey interaction by presenting a predator-infective prey model to analyse the dynamics and coexistence of the species in the community. After presenting the basic framework of the model we parametrize the model for an opportunistic pathogenic bacteria and the commercially cultivated sea cucumber, an economically important species in aquaculture. The sea cucumber is appreciated as a delicacy and aphrodisiac widely in Asia. Even though the catches from the wild populations have drastically declined, the production of cultured sea cucumbers in year 2014 was over 200000 tonnes in China alone. [16]. According our results the species most likely coexist at a stable equilibrium.

We also analysed the conditions for species extinctions. For the predator the extinction depends on the infectivity of the prey, and its population size as well as the attack rate of the predator. The possibility of recognizing the effects of an infective prey within a food web is thus meaningful both scientifically and economically.

### Conceptual model description

The predator (S) and the prey (C) interact according to a conventional predator-prey model (Fig 1). The predator population grows by consuming the prey (i). However, both species also use other resources for growth ((vi) and (vii)), meaning that each of them can survive as a single species population. Thus, we are dealing with a generalist predator. However, in a special case the predator can be specialist. As the prey is also pathogenic to the predator, a part of the predator population is infected (ii), increasing the population size of infected predators (I). An infected predator can recuperate (iii), die naturally (iv), or become a growth resource for the pathogenic prey (v).

**Fig 1.**
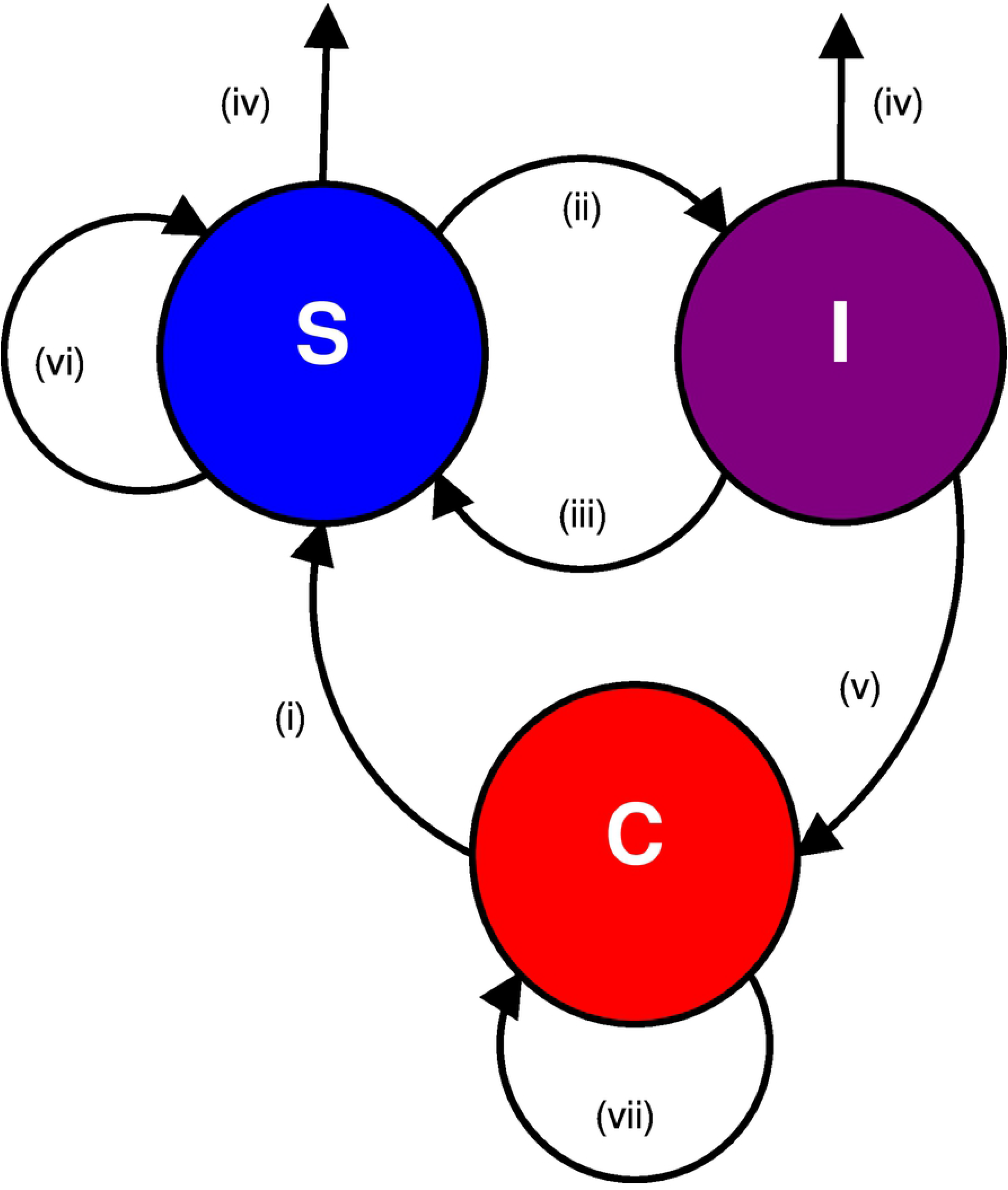
A schematic presentation of the predator-infective prey model. The predator (S) and the prey (C) interact according to a conventional predator-prey model. However, the prey is pathogenic to the predator, and a part of the predator population is infected (I). An infected predator can recover, die naturally, or become consumed by the pathogenic prey.

A distinctive aspect in our predator-prey interaction is that both the prey and the predator are only a part of a food web. Both species have a base growth rate that is independent of their mutual interaction, and they both are able to grow independently according to the respective carrying capacity of the environment ((vi) and (vii) in Fig 1).

### Modelling partial role reversal in predator-prey interaction

Let C, S and I denote the abundances of the prey, predator and infected predator populations, respectively. The differential equation model for the dynamics of the populations are given as

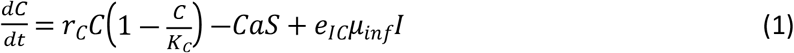

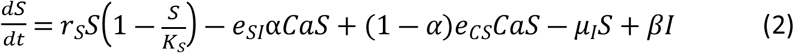

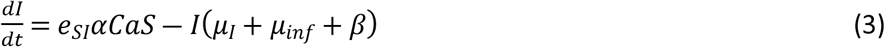

where the increases of the prey and predator abundances are both defined as logistic growth. Parameters *r_C_, K_C_, r_S_* and *K_S_* are the growth rates and carrying capacities of the prey and predator, respectively. Parameter *a* (0≤*a*≤1) denotes the attack rate of the predator *S*. This can be interpreted either as a fraction of the feeding area grazed during a time step, or it can equally be interpreted as the prey selectivity coefficient of the predator. Thus, the total number of the prey harvested by the predator is *aC*, and the harvesting is described as Type I functional response. Parameter *α* denotes the fraction of the infective prey from the total prey population. Thus, from the predation rate *CaS* fraction (1 – *α*) increases the growth rate of the healthy predator population with a prey to predator growth conversion efficiency *e*_CS_. The rest of the harvested prey, *αCaS*, infects the predators at a conversion rate e_SI_. The infected the predators end up in the infected predator population *I*. Parameter *β* denotes the recovery rate of the infected predators. Parameters *μ_inf_* and e_IC_ denote the predator infection mortality and predator to prey conversion efficiency, respectively. Finally, *μ_I_* denotes the natural predator mortality.

Our model follows the basic structure of the traditional predator-prey Lotka-Volterra model in that the predation is modelled as Type I functional response, and that the healthy harvest is used for the growth of the of the predator. The model differs from the Lotka-Volterra model in that both the susceptible predator and the infective prey have their own logistic growth functions implying that they are generalists rather than specialists. The infected predator population grows only at the expense of the infections of the healthy predators. The infected predators also serve as a resource for the prey growth as they become diseased.

### Parametrization of the model

Most of the parameters were chosen according to the suitable values found from the literature. Conversion efficiencies were calculated as the ratio of dry weights of the predator and the prey multiplied by ecological efficiency. Ecological efficiency was set to 0.25 for the sea cucumber, and to 0.5 for the bacteria [17,18].

Because we could not find the dry weight of *V. splendidus*, we used the dry weight of *E. coli* [19]. The dry weight of *A. japonicus* is calculated according to the article by Sun et al., where it was stated that the dry weight of *A. japonicus* equals 0.075×wet weight [4]. The mean wet weight *m_Aj_* was set at 150g [20].

Prey to predator conversion efficiency is calculated as 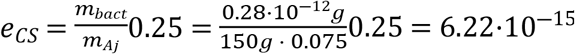.

Predator to prey conversion efficiency is 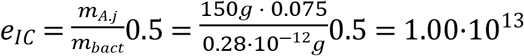. According to the empirical study by Lysenkov et al. the natural population density of *A. japonicus* is 0.14 individuals per square meter, even though the observed density has fallen to 0.023 individuals per square meter because of illegal harvesting [20]. Therefore, the area of the feeding unit is set to 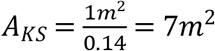 and depth of foraging to 1*cm*. We calculated the predator attack rate using the formula

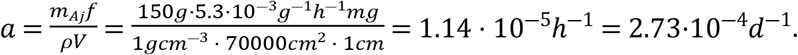

where *m_Aj_* is the wet weight of the sea cucumber and *f* is the amount of sediment eaten by the sea cucumber per hour per gram of sea cucumber [4], *ρ* is the density of the sediment as given by Kennish [9], and *V* is the volume of the feeding unit. The resulting attack rate is the nominal portion of the available prey eaten within a time step. Because the actual attack rate depends also from the selectivity of the predator, a range of attack rate values around the nominal value was used in model analysis and numerical simulations.

We have taken the carrying capacity of the bacteria *K_C_* from the literature [7,11]. For the sea cucumber carrying capacity *K_S_* we tested a range of values, but for consistency in the shown simulation results *K_S_* is always 10000.

The symbols and parameters used in the model and are shown in Table 1.

**Table 1.**
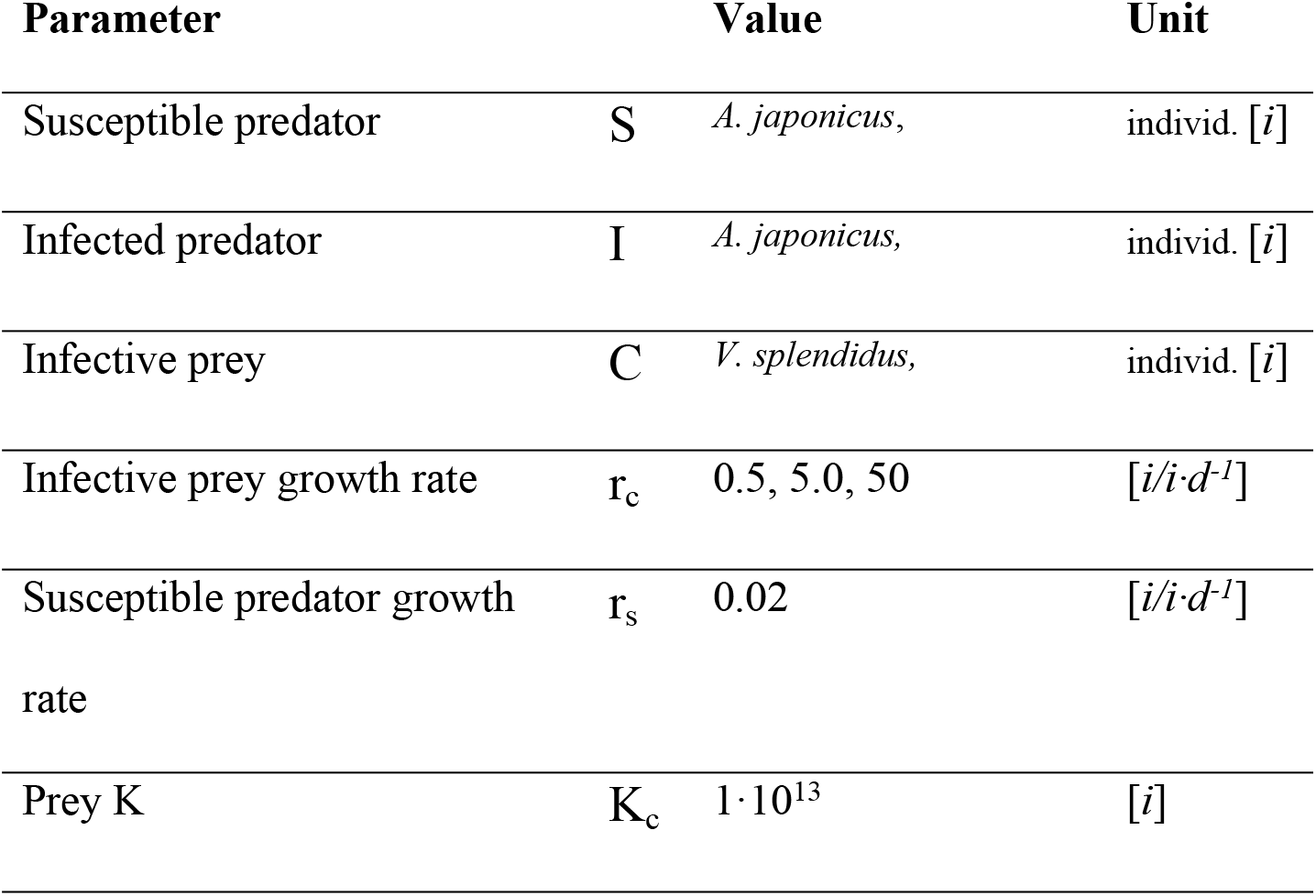
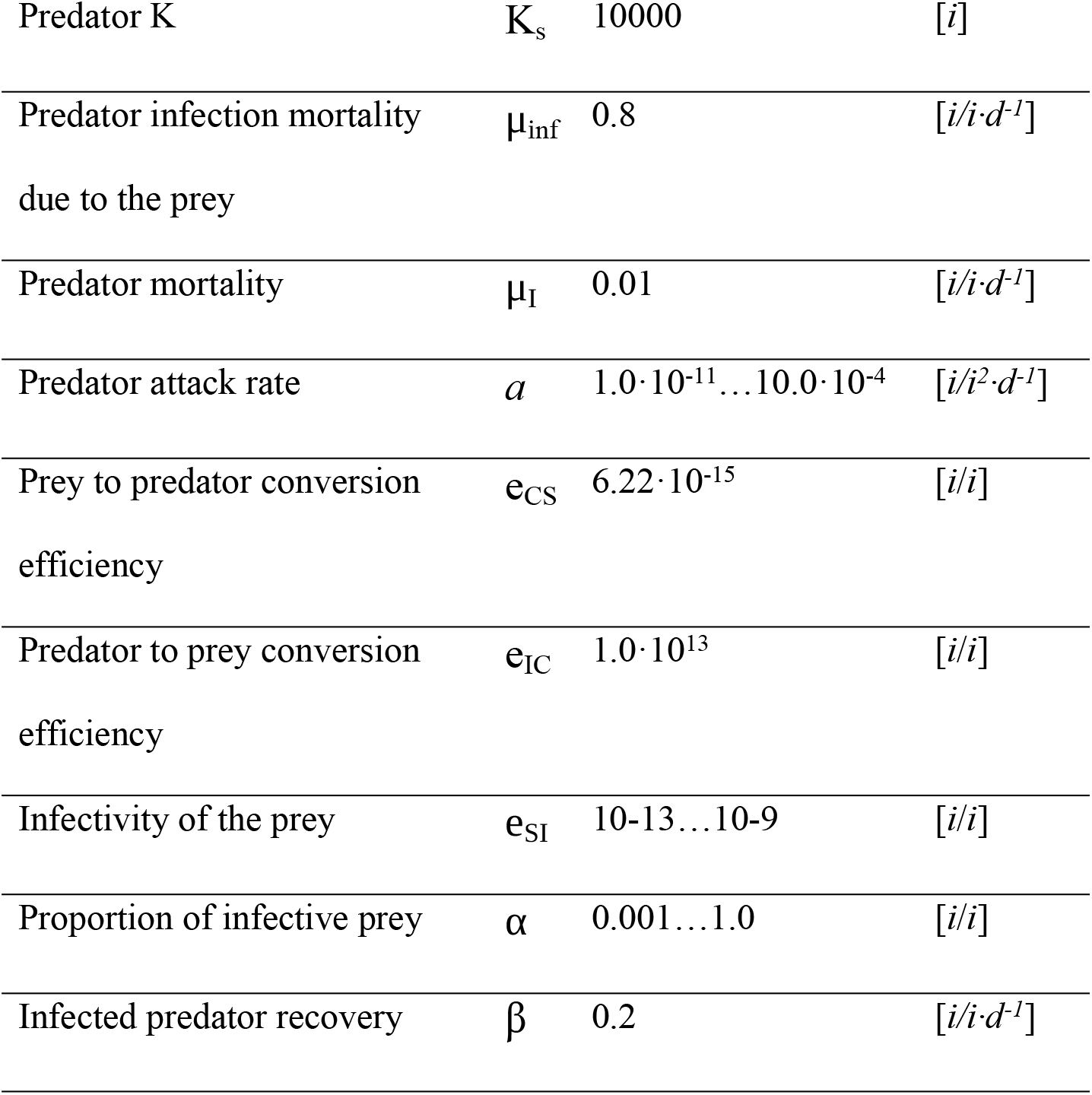
Symbols and parameter values

### Model analyses

#### Population equilibria

The equilibrium of the community is the starting point of the analysis of community behaviour. The equilibrium is defined by assuming the time derivatives in the population equations (1)-(3) equal to zero:

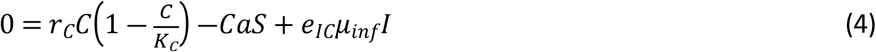

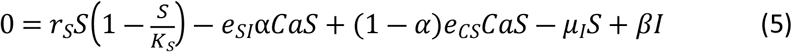

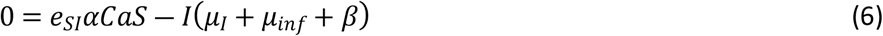

Inserting *I* from eq. (6) into (4) and (5) and dividing the resulting equations by C and S, respectively, we get

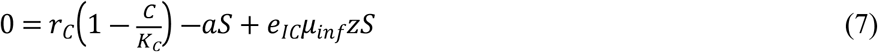

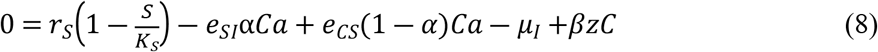

where 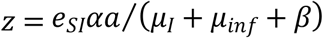

Linear equations (7) and (8) can be presented in a matrix form

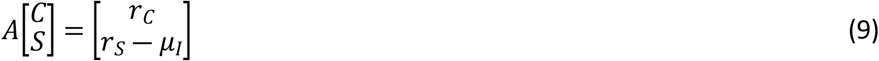

where

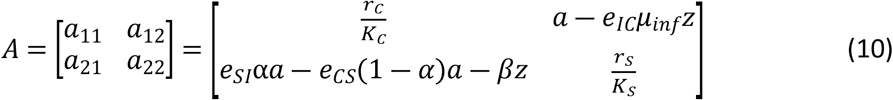

The solution of eqs. (9) and (10) is given as

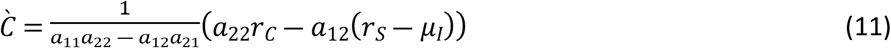

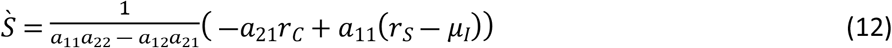

Infected predators are then calculated as 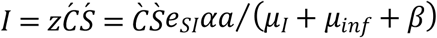 (eq. (6)).

The equilibrium states of interest are the following:

a. Both species coexists at a general equilibrium: 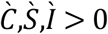. Stability of this equilibrium represent continuing coexistence of the species.
In the absence of species interaction the environmental carrying capacity of the prey is equal to KC and that of the predator is equal to (*r_S_* – *μ_I_*)K_S_/r_S_.
b. Infective prey exists but is zero: 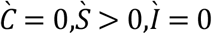. This represent an extinction of the prey.
c. Predator is absent: 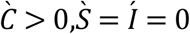. This represents an extinction of the predator.

### Analytical results

The analysis of the model presents us three possible outcomes. Either the prey or the predator drives the other to extinction, or both populations coexist in a stable equilibrium.

### Numerical simulations

The numerical simulations of the model (1)-(3) were performed out using Matlab R2020b. Numerical simulations were in accordance with the analytical results (Section “Population equilibria”). Both the prey and the predator were able to drive the other to extinction. All simulation results with positive coexisting populations were locally stable. The parameters used in the simulations are described in Section “Parametrization of the model”. Because simulations exemplify the partial role reversal between *A. japonicus* and *V. splendidus*, respective conversion efficiencies were used throughout, as well as the high mortality rates shown to be associated with the infection [12,13]. The effects of infectivity of the prey e_SI_, proportion of the infective prey α, and the attack rate *a* were tested using wide parameter ranges. Though the outbreaks caused by *V. splendidus* have been associated with high mortality rates, we tested the model also with low infection mortality rates and high recovery rates. Even when infection mortality μ_inf_ and infected predator recovery β were 0.3 and 0.7, respectively, the results remained qualitatively same. Initial population sizes did not affect the results, and the model gives consistent results.

## Results

For an opportunist prey with a high environmental growth rate the level of infectivity, e_SI_, is not crucial (Fig 2). The prey population size will settle around the level of carrying capacity K_C_. A low infectivity e_SI_ combined with a high environmental growth rate r_C_ of the prey can be beneficial also for the predator because the predator is able to sustain population levels above the environmental carrying capacity (*r_s_ – μ_I_*)*K_S_/ r_S_*. Rising the level of infectivity, however, decreases the predator population. To the contrary, the population size of a prey with slow environmental growth depends on the level of infectivity. Low infectivity leads to the extinction of the slowly growing prey.

**Fig 2.**
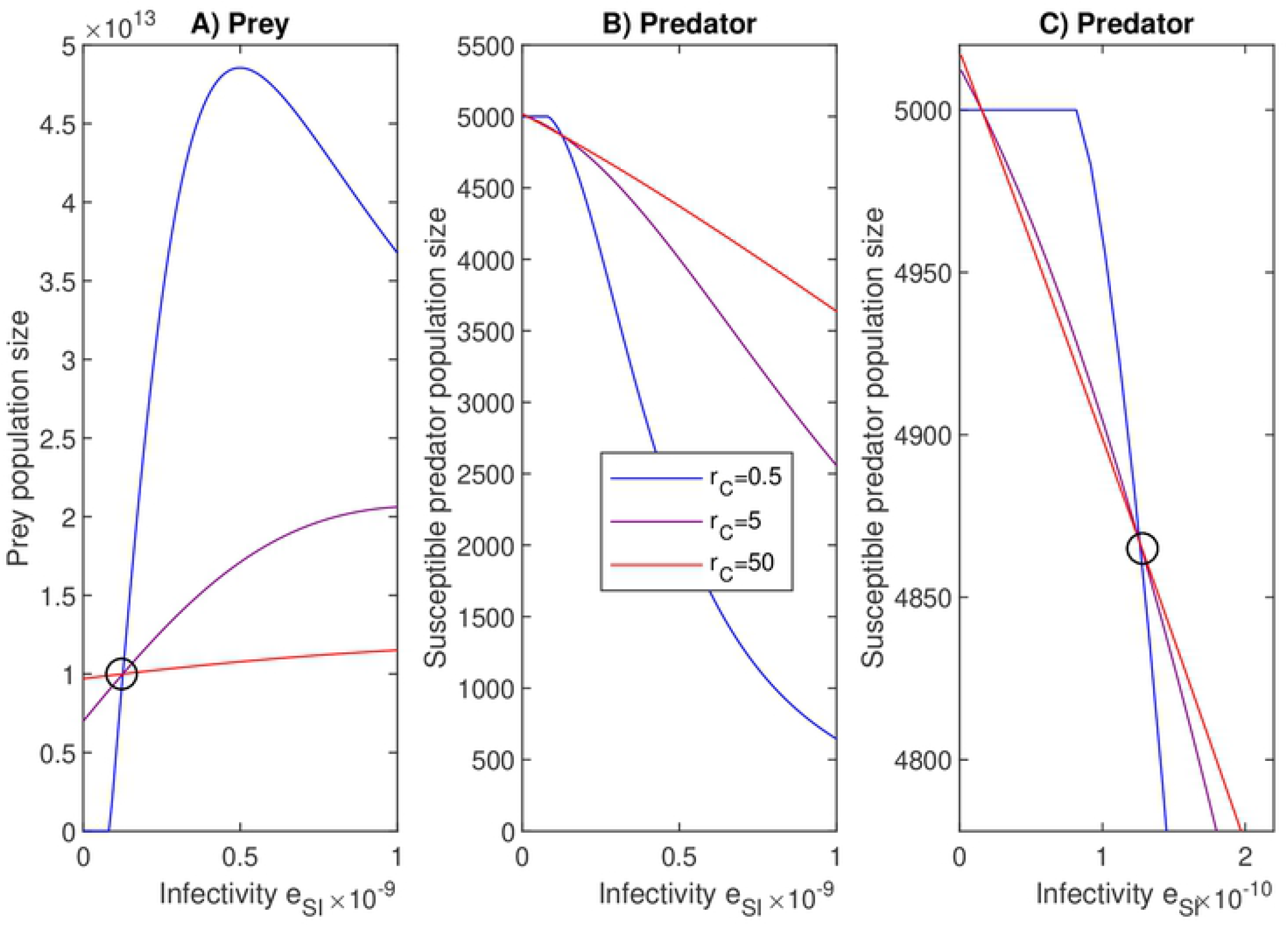
Infectivity affects both prey and predator population sizes. For low infectivity e_SI_ a higher outside prey growth rate *r_C_* supports larger prey population than a lower growth rate, but this is reversed if e_SI_ increases enough. After the turning point (o), where high and low growth rates of the prey provide equal population sizes, an increase in infectivity e_SI_ of the prey results in greater prey and lesser predator population sizes. If the infectivity is increased even more, the trend of the prey population turns into decreasing. At the extinction of the prey (at low infectivity values and low prey growth rate) the predator population size settles down at its environmental carrying capacity 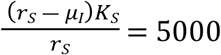. Subfigures A and B show the full scale of the population sizes, whereas subfigure C displays a closer view to the predator population at the turning point (o). Red, purple and blue lines are fast (r_C_=50), medium (r_C_=5) and slow (r_C_=0.5) growth rates. The predator’s attack rate *a=3.0·10^−4^* and the infective proportion of the prey α=0.001. Infectivity e_SI_ ranges from 10^−13^ to 10^−9^.

High infectivity e_SI_ increases the population size of a slowly growing prey because increasing infectivity allows the prey to reach a higher prey population size as compared to a prey with a higher growth rate r_C_ (Fig 2). However, due to the high mortality μ_inf_ associated with the infection, a too high level of infectivity causes the extinction of the predator and a decline in the prey population size. Likewise, in the case of fast growing prey, very high infectivity leads the prey population size to settle at the environmental carrying capacity.

Fig 2 also illustrates the presence of a turning point such that the order of the population sizes will change with the change of a parameter. When e_SI_ = 1.22e-10, making *a_12_ =* 0 (eq. 8), the equilibrium population size of the prey equals to its environmental carrying capacity 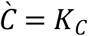. At the same value of infectivity the equilibrium population size of the predator will be 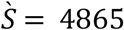. Below the turning point slow prey growth rates supports lower prey population sizes than higher growth rates. When the parameter e_SI_ passes the turning point then the order of the population sizes is reversed. The effect of the turning point to the population sizes of the predator is opposite. Note that the turning point is not a uniquely defined concept but always related a chosen parameter. This is because the condition α_12_ = 0 can become true for choosing appropriate values for *e_SI_,α, e_IC_* and *μ_inf_*. A comparable analysis can be carried out for the solutions with α_21_ = 0.

Infectivity e_SI_ and the proportion of infective prey in the total prey population α have parallel but not completely interchangeable effects on the population sizes of the prey and predator (Fig 3). If the prey is very weakly infective (e_SI_=10^−13^, Figs 3A,B), the predator will survive any proportion of the infective prey, and can even completely eradicate a slow growing prey. In contrast, if the infectivity is high (e_SI_=10^−11^, Figs 3C,D), then the predator will become extinct even at relatively low infective prey densities. This happens regardless of the prey growth rate.

**Fig 3.**
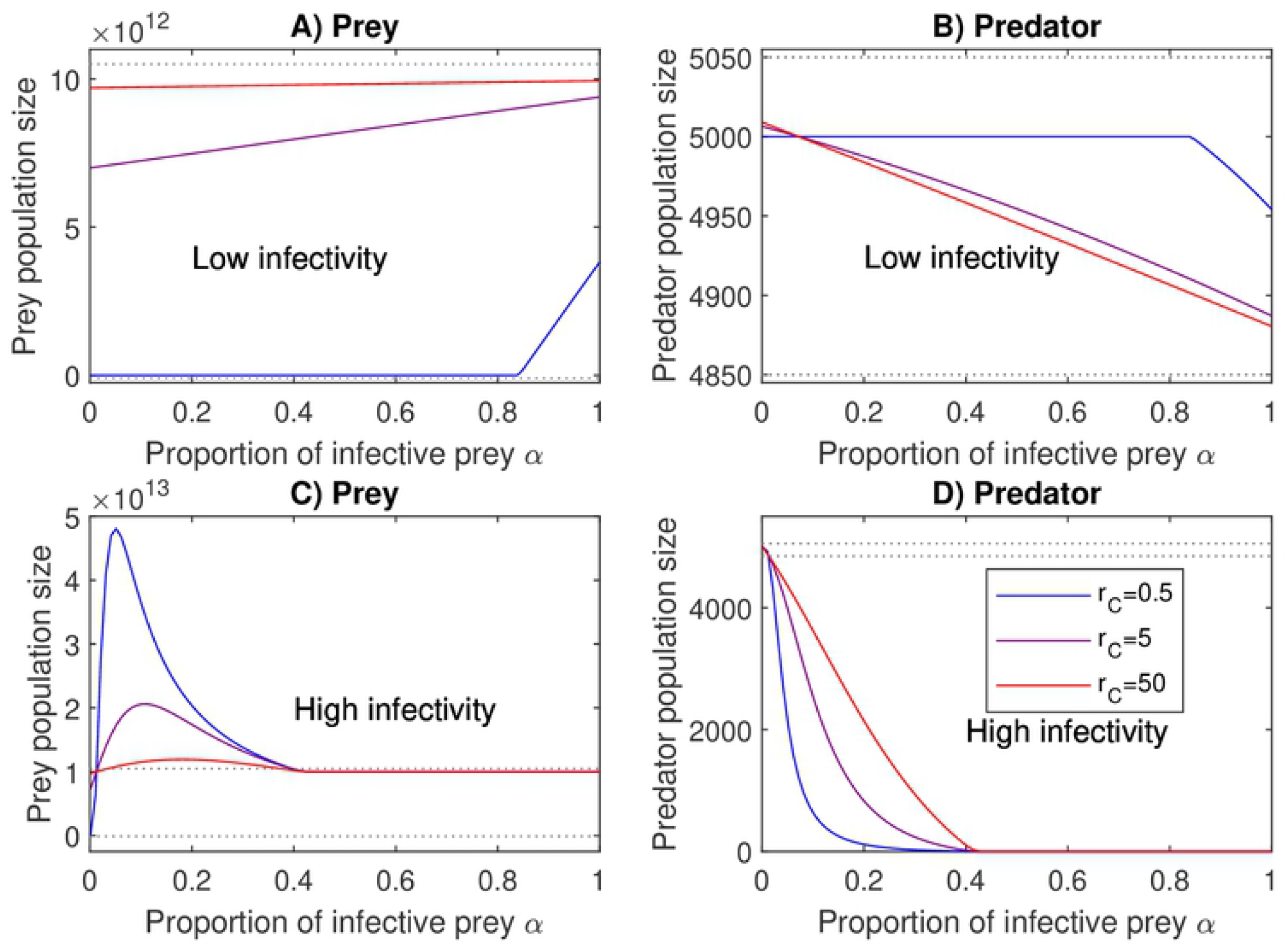
Proportion of infective prey *α* affects the population in the same way as infectivity *e*_SI_. Low infectivity of prey, e_SI_, supports lower prey population sizes (A) and higher predator population sizes (B) than high infectivity (C and D, respectively). Slowly growing prey with low infectivity can proliferate only if the majority of the prey are infective (A). Even high proportions of infective prey cause only a slight decrease in predator population (B). High infectivity e_SI_ prevents the extinction of the prey. Highly infective prey thrives best when it forms relatively small part of the prey population (C) because the predator becomes extinct if the majority of the prey are infective (*α* ≳ 0.4) (D). The whole range of final population sizes in subfigures A and B fit within the dotted lines in subfigures C and D, respectively. Red, purple and blue lines are fast (r_C_=50), medium (r_C_=5) and slow (r_C_=0.5) growth rates. Infectivity values are e_SI_=10^−13^ in subfigures A and B, and e_SI_=10^−11^ in subfigures C and D.

If attack rate approaches zero both the prey and the predator population sizes tend towards the environmental carrying capacity regardless of the infectivity (Fig 4). Increasing attack rate may have different effects on the prey and predator sizes. When the value of the infectivity remains low an increase in the attack rate benefits the predator (Fig 4B). High growth rate of the prey results in larger predator population than low growth rate. If the value of the infectivity is increased slightly (moderate infectivity) an increment in growth rate decreases predator population sizes (Fig 4D). In both cases increasing attack rate decreases the prey population size (Figs 4A,C).

**Fig 4.**
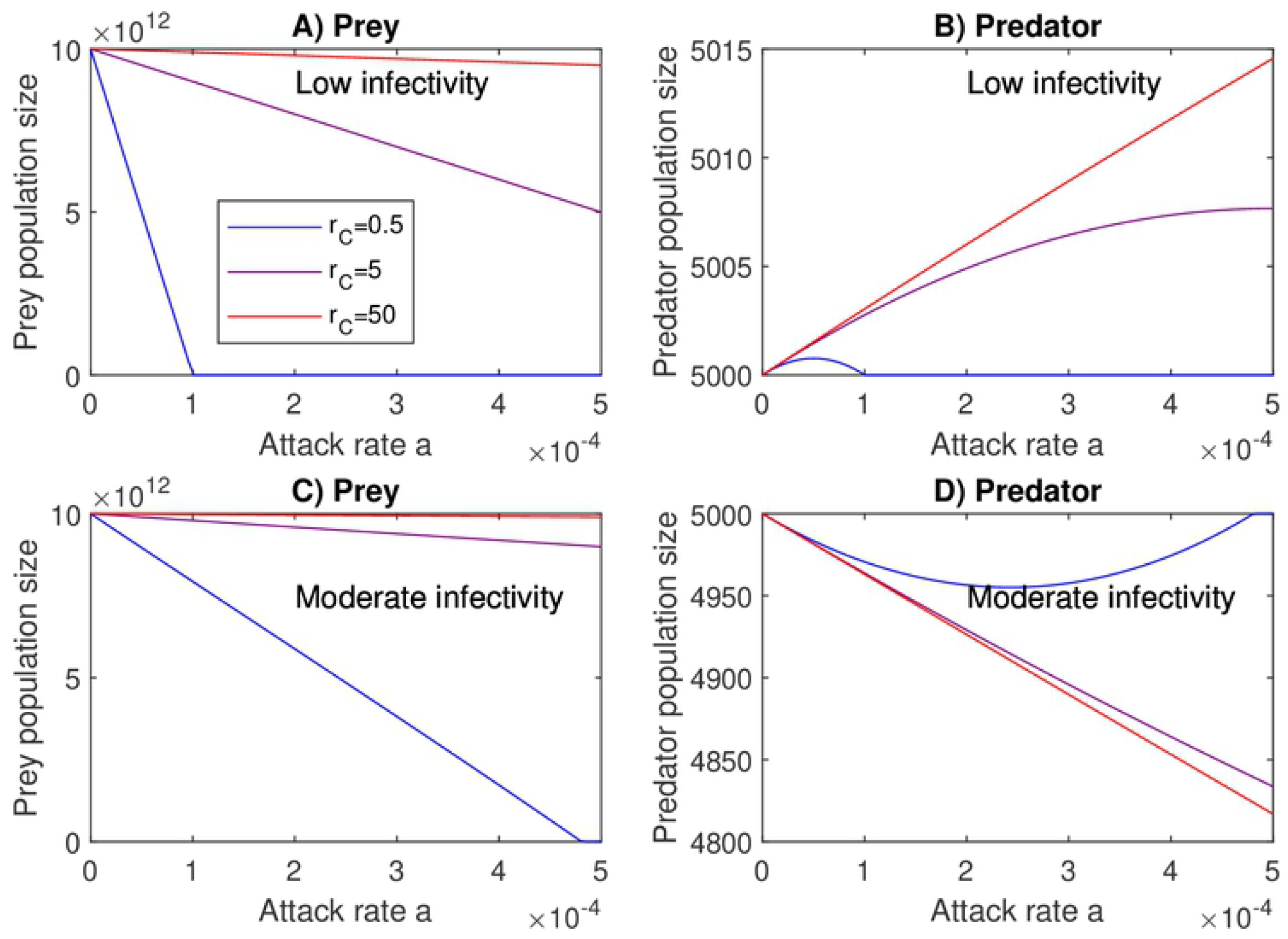
An increase in the infectivity may reverse the effect of predator attack rate on the predator population size. In subfigures A and B the prey’s infectivity *e_SI_* is weak. Increasing the predator attack rate *a* decreases prey and increases predator population sizes. In contrast, *e*_SI_ in subfigures C and D is slightly greater, and increasing attack rate decreases the population levels of the prey as well as of the predator. As the attack rate decreases, the population sizes approach their respective environmental carrying capacities. Red, purple and blue lines are fast (r_C_=50), medium (r_C_=5) and slow (r_C_=0.5) growth rates. The infectivity values are e_SI_=10^−13^ in subfigures A and B, and e_SI_=10^−10^ in subfigures C and D.

### Extinction of the species

We consider here the possibility of extinction of the predator or the prey. The questions of interest are: 1) Under which conditions the predator can drive the prey to extinction such that the species community would approach lie at a “predator only” equilibrium 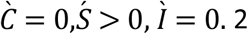) Alternatively, we ask under what conditions the prey can eradicate the predator such that the species community would ultimately lie the “prey only” equilibrium 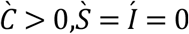.

Consider first the “predator onl” equilibrium 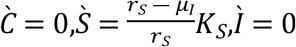, that is, the sea cucumber lies at its carrying capacity and *V. splendidus* has been driven to extinction. If the equilibrium is locally stable then the extinction of *V. splendidus* is expected to occur. The local stability of the linearized dynamics at the equilibrium can be analysed studying the properties of the following Jacobian matrix

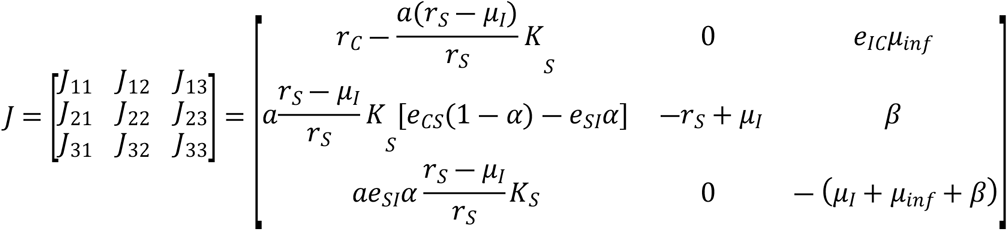

Recall that if the real parts of the eigenvalues of J are all negative the system is locally stable. It can be shown that the first eigenvalue is λ_1_= — *r_s_* + *μ_I_* which we assume to be negative. The remaining two eigenvalues depend on the submatrix where line 2 and column 2 are deleted in matrix *J*. The eigenvalues *λ_2_,λ_3_* both have negative real pars if and only if [21]

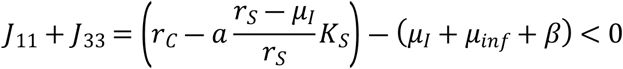

and

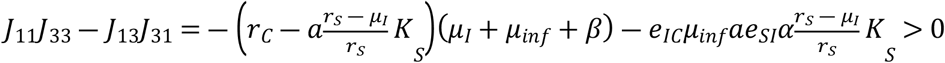

In this case extinction occurs. For example, if the proportion of infective prey is low (*α* ≈ 0) and the growth rate is low 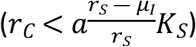 then the both conditions become true and the predator will eradicate the prey. High proportion of infective prey, high energetic efficiency and high carrying capacity may protect the prey from extinction.

The extinction of the prey depends crucially also on the attack rate of the predator (Fig 5). There is a threshold value or a minimum attack rate at which the predator can cause the extinction of the prey.

**Fig 5.**
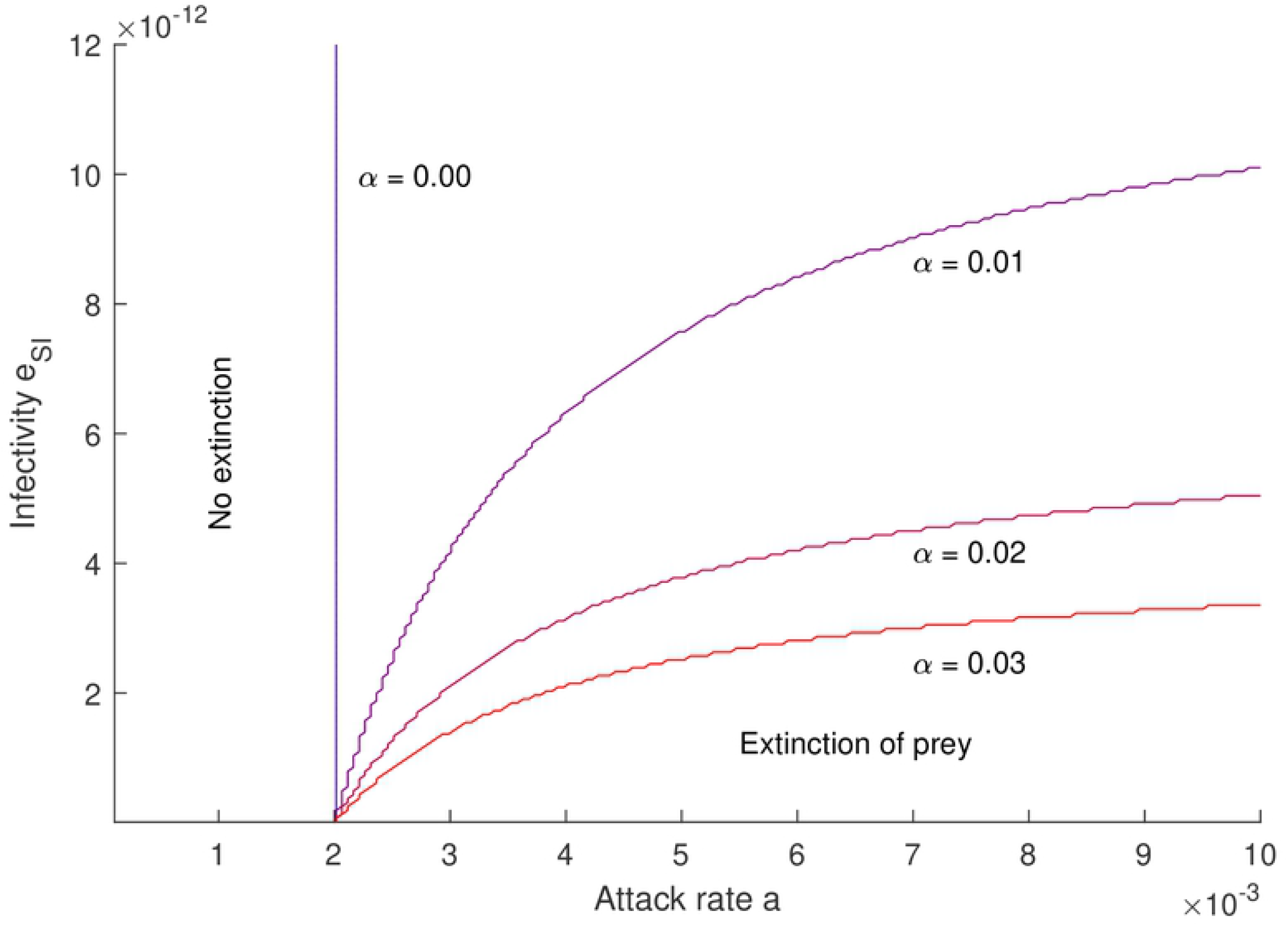
Predator attack rate affects the extinction of prey. Extinction of prey is possible if the predator’s attack rate is higher than the threshold defined by the prey’s growth rate. Fast growing prey survives higher predator attack rate than slow growing. If the prey’s infectivity e_SI_ is high, only a small fraction α of the prey population needs to be infective to escape the extinction. Prey growth rate r_C_=10.

If the prey is a specialist such that it consumes only the predator (*r*_C_=0), then the extinction can be due to low infectivity or insufficient infective population. Yet, even a low prey growth rate may keep the prey population alive, if the prey is infective enough. Because infective prey can survive when its outside growth rate *r_C_*=0, a positive growth rate guarantees the survival also in the case of relatively low infectivity, if the infective prey forms large part of the predator’s diet.

Extinction of the predator leads to the “prey only” equilibrium 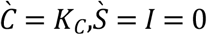. The linearized dynamics of the predator-prey interaction at the equilibrium can be presented as

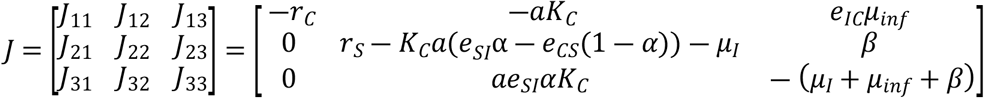

The first eigenvalue of the Jacobian matrix J is *Λ*_1_= –*r_C_*. The remaining two eigenvalues depend on the submatrix with line 1 and column 1 deleted in the Jacobian matrix *J*. The real parts of the eigenvalues *λ_2_,λ_3_* are both negative if and only if [21]

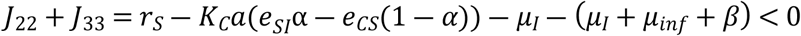

and

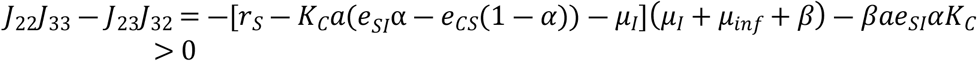

The proportion of the infective prey α is crucial. Assume that α =0. Then J_22_J_33_ – J_23_ J_32_ = – [r_S_ + K_C_a – μ_I_], μ_I_ + μ_inf_ + β)<0 indicating that the predator does not become extinct. Assume next that α =1. Then *J*_22_ + *J*_33_ < 0 if

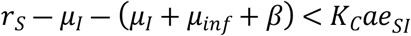

and *J*_22_*J*_33_ – *J*_23_*J*_32_ > 0 if – *K_C_ae_SI_*(*μ_I_* + *μ_inf_*) – (*r_S_* – *μ_I_*)(*μ_I_* + *μ_inf_* + *β*) > 0, indicating that the predator will become extinct.

Under these conditions the predator becomes extinct. Fig 6 presents how the change in infectivity e_SI_ together with the different infective prey proportions affect the extinction of predator. If the prey’s growth rate r_C_ is at all positive, the prey can drive the predator to extinction. Interestingly, if r_C_>0 the level of growth rate does not affect the results.

**Fig 6.**
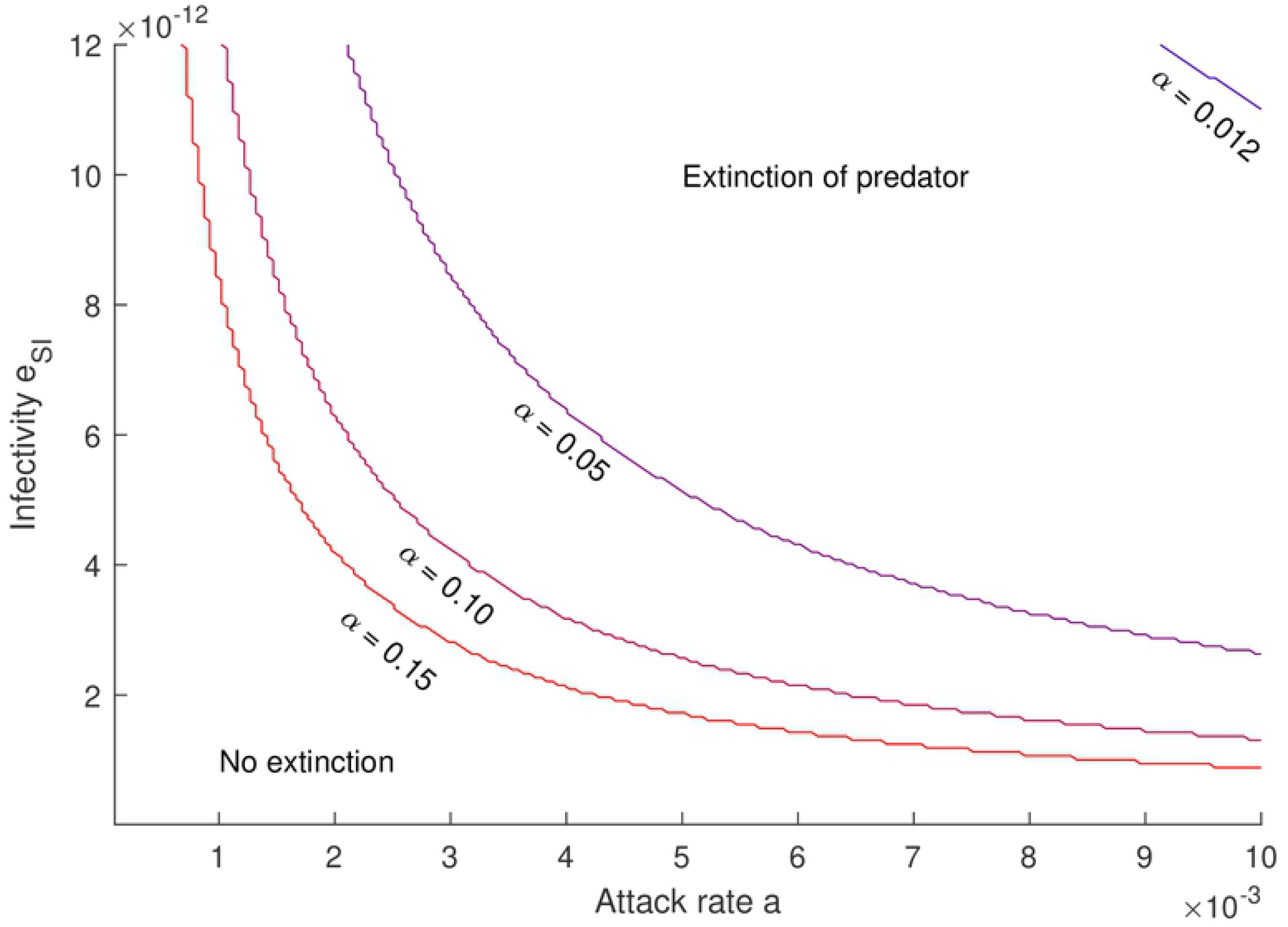
Extinction of predator depends on the volume of infective prey consumption. If the infective prey forms a large fraction of the available prey population, lower prey infectivity e_SI_ and predator attack rate are needed to eradicate the predator. To the contrary, if the infective fraction α is small, or the infectivity e_SI_ is low, the predator prevails. Using the parameters for *A. japonicus* and *V. splendidus* the predator survives always if α<0.011. At very low attack rate or infectivity the predator does not consume enough of the infective prey to suffer extinction. In the case of a specialist prey, when r_C_=0, it can not drive the predator to extinction. However, as long as r_C_>0, then the extinction does not depend on the rate of growth.

## Discussion and conclusions

We have presented a new predator-prey model with partial role reversal where the predator can become a target of attacks by the prey such that the prey can use the predator as a resource for growth. We parametrized the model using sea cucumber *A. japonicus* and a bacterium species *V. splendidus* as a model system.

A distinctive feature in our model is that both the prey and the predator are only a part of a food web. Both species have an environmental growth rate that is independent of their mutual interaction, and they both are able individually to grow to their respective carrying capacity. Thus, both the prey and the predator are generalists. The predator can benefit from the infective prey through eating the increasing prey population. The final size of the predator population depends on the proportion of infective prey in its diet. A small proportion of very infective prey functions similarly with a large proportion of less infective prey.

Overall, the partial role reversal in the predator prey community stabilizes the predator-prey interaction. The same effect is observed if we add logistic growth to both species in the Lotka-Volterra equation. We did not observe in our extensive simulations any signs of instability in the coexistence solutions

We also analysed the conditions for species extinction. A generalist predator becomes eradicated if some part of the prey population is infective, the infection can cause mortality, and the abundance of the infective prey is high. For a specialist predator the extinction depends on the infectivity of the prey, and its population size as well as the attack rate of the predator. The extinction of the prey is possible if its infectivity is low and either it the infective prey forms only a small fraction of total prey population or the growth rate is low.

The infective prey-predator model departs from a predator-non infective prey model by quantifying the infectivity of the prey and the mortality of the infection. In principle, this resembles a fatal infectious disease. However, the predator is also able to consume the prey regardless of its pathogenicity, and can therefore benefit from the growing pathogenic prey population. It is noteworthy that infection does not need to be a bacterial infection. Any similar situation can play the part of a disease in the model framework. The examples include a shoal of young pikes can that attract a growing number of sticklebacks [3]. In these examples, the infectivity is taken as a number that describes the ability of the prey to find potential victims among the predators and to attract the rest of the prey population to the site. The infection mortality describes the probability of death of an infected or attacked predator.

The system consisting of a predator and an infective prey remains mostly an unresearched subject. The population model presented here describes the process of role reversal using several parameters. However, of these parameters only a few were in the model markedly involved in the role reversal process. These include the prey to predator and predator to prey conversion efficiencies, and the infectivity of the prey. This implies that it would be possible to address the subject empirically by studying a suitable pair of model organisms.

Aquaculture provides many opportunities to find both scientifically and economically interesting targets for basic and applies research. Also, agriculture can be considered as a field that would benefit from the research.

For the purposes of enhancing sea cucumber cultivation, factorial experiments manipulating the growth conditions of the non-pathogenic vs. pathogenic bacteria as food could be set up. One avenue on this would be for example a biological control [22] of *Vibrio* using specific lytic bacteriophages continuously of periodically added to the culturing ponds. Bacteriophages are commonly species specific and can be mass-produced in bioreactors. In practise, *Vibrio* phages could be isolated from raw water samples by filtering out the bacteria and adding the filtrate on *Vibrio* pure culture to amplify only *Vibrio*-specific phages.

## Acknowledgements

We thank Raine Kortet for useful discussions.

## References

1. Barkai A, McQuaid C. Predator-prey role reversal in a marine benthic ecosystem. Science 1988; 242(4875): 62–64.

2. Sánchez-Garduño F, Miramontes P, Marquez-Lago T. Role reversal in a predator-prey interaction. R Soc Open Sci 2014;1:140186.

3. Nilsson J, Flink H, Tibblin P. Predator–prey role reversal may impair the recovery of declining pike populations. Journal of Animal Ecology 2019; 88 (6): 927–939.

4. Sun J, Zhang L, Pan Y, Lin C, Wang F, Kan R, et al. Feeding behavior and digestive physiology in sea cucumber *Apostichopus japonicus*. Physiology & Behavior 2015; 139: 336–343.

5. Xu Q, Hamel J-F, Mercier A. Chapter 10 - Feeding, digestion, nutritional physiology, and bioenergetics. Developments in Aquaculture and Fisheries Science 2015; 39: 153–175.

6. Bratbak G. Bacterial biovolume and biomass estimations. Applied and Environmental Microbiology 1985; 49(6),:1488–1493.

7. Watson SW, Novitsky TJ, Quinby HL, Valois FW. Determination of bacterial number and biomass in the marine environment. Applied and Environmental Microbiology 1977; 33(4): 940–946.

8. Liu X, Zhou Y, Yang H, Ru S. Eelgrass detritus as a food source for the sea cucumber *Apostichopus japonicus* Selenka (Echinodermata: Holothuroidea) in coastal waters of north China: An experimental study in flow-through systems. PLOS ONE 2013; 8(3): e58293

9. Kennish MJ. Encyclopedia of estuaries. 2016, Springer Netherlands, ISBN 978-94-017-8800-7.

10. Xu H, Wang L, Bao X, Jiang N, Yang X, Hao Z, et al. Microbial communities in sea cucumber *(Apostichopus japonicus)* culture pond and the effects of environmental factors. Aquaculture Research 2019; 50: 1257–1268.

11. Vezzulli L, Pezzati E, Stauder M, Stagnaro L, Venier P, Pruzzo, C. Aquatic ecology of the oyster pathogens *Vibrio splendidus* and *Vibrio aestuarianus*. Environmental Microbiology 2015; 17(4): 1065–1080.

12. Deng H, He C, Zhou Z, Liu C, Tan K, Wang N, et al. Isolation and pathogenicity of pathogens from skin ulceration disease and viscera ejection syndrome of the sea cucumber *Apostichopus japonicas*. Aquaculture 2009; 287 (1-2): 18–27.

13. Liu N, Zhang S, Zhang W, Li C. Vibrio sp. 33 a potential antagonist of *Vibrio splendidus* pathogenic to sea cucumber (*Apostichopus japonicus*). Aquaculture 2017; 470: 68–73.

14. Zhang Z, Lv Z, Zhang W, Shao Y, Zhao X, Guo M, et al. Comparative analysis of midgut bacterial community under *Vibrio Splendidus* infection in *Apostichopus japonicus* with hindgut as a reference. Aquaculture 2019a; 513: 734427.

15. Zhang Z, Zhang W, Hu Z, Li C, Shao Y, Zhao X, et al. Environmental factors promote pathogen-induced skin ulceration syndrome outbreak by readjusting the hindgut microbiome of *Apostichopus japonicus*. Aquaculture 2019b; 507: 155–163.

16. Han Q, Keesing JK, Liu D. A Review of sea cucumber cquaculture, ranching, and stock enhancement in China. Reviews in Fisheries Science & Aquaculture 2016; 24(4): 326–341.

17. Yang H, Yuan X, Zhou Y, Mao Y, Zhang T, Liu Y. Effects of body size and water temperature on food consumption and growth in the sea cucumber *Apostichopus japonicus* (Selenka) with special reference to aestivation. Aquaculture Research 2005; 36: 1085–1092.

18. Yingst JY. The utilization of organic matter in shallow marine sediments by an epibentic deposit-feeding holothurian J Exp Mar Biol Ecol 1976; 23: 55–69.

19. Neidhardt FC, Curtiss IR, Ingraham JL, Lin, ECC, Low KB, Magasanic B, et al., editors. Eschericia coli and salmonella: Cellular and molecular biology. 2^nd^ ed. Washington D.C.: ASM Press; 1996.

20. Lysenko VN, Zharikov VV, Lebedev AM. The current status of populations of the sea cucumber *Apostichopus japonicus* (Selenka, 1867) in the Far Eastern marine reserve. Russian Journal of Marine Biology 2018; 44(2): 164–171.

21. Edelstein-Keshet, L. Mathematical models in biology. 1^st^ ed, Random House, NY, 1988.

22. Merikanto I, Laakso JT, Kaitala V. Outside-host phage therapy as a biological control against environmental infectious diseases. Theoretical Biology and Medical Modelling 2018; 15(7).

